# Neonicotinoids disrupt memory, circadian behaviour and sleep

**DOI:** 10.1101/2020.04.07.030171

**Authors:** Kiah Tasman, Sergio Hidalgo, Edgar Buhl, Sean A. Rands, James J.L. Hodge

## Abstract

Globally, neonicotinoids are still the most used insecticides, despite their well-documented sub-lethal effects on beneficial insects^1^. Neonicotinoids are agonists at the nicotinic acetylcholine receptors, the main mediator of synaptic transmission in the insect brain^2–5^, making them highly potent neurotoxins and insecticides^6,7^. Memory, circadian rhythmicity and sleep are essential for efficient foraging in many pollinating insects, and involve nicotinic acetylcholine receptor signalling^2,4,8–10^. The effect of field-relevant concentrations of European Union-banned neonicotinoids: imidacloprid, clothianidin and thiamethoxam, as well as the currently unbanned thiacloprid were tested on *Drosophila* memory, circadian rhythms and sleep. Field-relevant concentrations of imidacloprid, clothianidin and thiamethoxam disrupted learning, behavioural rhythmicity and sleep whilst thiacloprid exposure only affected sleep. Exposure to imidacloprid and clothianidin directly affected neurophysiology, preventing the day/night remodelling and accumulation of pigment dispersing factor neuropeptide in the dorsal terminals of clock neurons. Knockdown of the neonicotinoid susceptible Dα1 and Dβ2 nicotinic acetylcholine receptor subunits in the mushroom bodies or clock neurons recapitulated the neonicotinoid like deficits in memory or circadian/sleep behaviour demonstrating that neonicotinoid effects are likely mediated in the mushroom body and clock circuitry. Disruption to learning, circadian rhythmicity and sleep are likely to have far-reaching detrimental effects on beneficial insects in the field.

## Introduction

An estimated 84% of European crops are dependent on pollinators whose service is valued at >€22bn/year and is essential to food security^11,12^. However, populations of pollinating insects are declining dramatically. For instance flying insects have decreased by over 75% in Germany over the last 27 years^13^. Diminishing pollinator numbers are a serious threat to our food security^12,14^, with intensive use of insecticides being implicated in these losses^12,14^. However, a third of the global crop is lost to pests and without pesticides this loss could be 75%, keeping the demand for insecticides high^15,16^. The most common insecticides worldwide are neonicotinoids, which account for 24% of the global insecticide market valued at $1 billion/year^16,17^. Neonicotinoids are highly efficacious non-specific neurotoxins, affecting both target pest species such as aphids and non-target beneficial insects. They share a mechanism of action, being agonists of nicotinic acetylcholine receptors (nAChR), the main neurotransmitter in the insect nervous system. They also display target site cross-resistance in pests, diminishing their effectiveness as insecticides and unfortunately encouraging application of increasing concentrations^7,17^. They were branded safe compared to previous insecticides because they do not act on mammalian nAChRs^7,17^. However, few precursive safety tests were performed on beneficial insects, for which neonicotinoids are now known to be potent neurotoxins with well-documented lethal and sub-lethal effects^11,15,17,18,19^. Therefore, continued intensive use is likely to have severe consequences on insect species numbers, with knock-on effects for the ecosystem, aquatic life, birds and mammals in addition to potential toxicity to humans^11,20–22^. Despite the recent European Union (EU) ban of three major neonicotinoids (the nitroimines: imidacloprid, clothianidin and thiamethoxam, the latter being a prodrug for clothiandin^23^), neonicotinoids remain the most widely used class of insecticide globally, with a number of studies showing there has been no decrease in the quantity of banned neonicotinoids found in different populations of honey and bumble bee across Europe a year after the ban^24,25^. Furthermore, some national governments have granted multiple exemptions for the spraying of oil seed rape and a number of other applications^26^, and neonicotinoids have high solubility and persistence in the environment^11^. Additionally, the cyanoimine neonicotinoid, thiacloprid is not currently covered by the EU ban, although its current status will be reappraised during 2020^27^ making findings on the sub-lethal effects of thiacloprid of outstanding importance. Therefore, despite the current EU ban, insects are still at risk of neonicotinoid exposure.

In the field, concentrations of neonicotinoids encountered by non-target insects are typically between 1-51 μg/L for seed treated crops and 61-127 μg/L for sprayed crops^19^. Concentrations as low as 1 µg/L (or 1 part per billion (ppb)) can cause significant behavioural effects due to the high potency of neonicotinoids, such as reduced foraging motivation in the bumblebee *Bombus terrestris*^28^. The potential sub-lethal effects of neonicotinoids are very far-reaching because of the central role of nAChR in synaptic neurotransmission in the insect brain^7,17^. Neonicotinoids cause this ligand-gated ion channel to open, thereby depolarising the neuron and increasing excitability. Prolonged exposure to the depolarising agonist may result in depolarising block, through voltage inactivation of voltage-sensitive Na^+^ channels required for action potential firing and nAChR desensitization^7,29^. This effect could have pronounced effects on memory formation and consolidation, which are critical for effective foraging in many pollinating insects.

Previous research in *Drosophila* demonstrated both the Kenyon cells, which constitute the insect memory centre called the mushroom body^5^, and their output neurons^2^, which mediate memory valence, are nicotinic with both brain regions also regulating sleep^10^. In honeybees, sub-lethal neonicotinoids electrically inactivated^29^ and decreased synaptic density^30^ of mushroom body neurons and resulted in disrupted olfactory memory^29^. Neonicotinoids also reduced honeybee antennal lobe Ca^2+^ responses and caused sensory deficits^31^, potentially indirectly causing olfactory memory deficits.

Memory formation is also reliant on circadian rhythms^32^ and sleep^8,33^. The effect of neonicotinoids on circadian rhythms and sleep is unknown. However work in *Drosophila* has shown that the setting of the central clock, synchronicity within the clock and communication between the light sensing organs and the central clock requires nAChRs signalling^3,4,34,35^. The timing of sleep/wake cycles is also determined by the circadian clock36 with the key clock neurons that mediate arousal and sleep being nicotinic^4,35^.

The pacemaker neurons of the insect clock consist of the pigment dispersing factor (PDF) neuropeptide expressing small and large ventral lateral neurons (s- and l-LNvs). The s-LNvs maintain rhythmicity in constant conditions and set the pace of the insect clock via PDF signalling^37^. The LNvs are nicotinic^3,4^ receiving ACh from the visual circuit including the lamina, with the s-LNvs also receiving ACh-mediated light input information from the Hofbauer-Buchner (HB) eyelets. These excitatory signals regulate the electrical excitability of the LNvs required for circadian function^38^. The LNvs are more depolarised and have an increased firing rate in the day than at night^39^. These day/night differences in excitability help sustain the molecular oscillation of clock genes in constant conditions as well as regulating s-LNv terminal remodelling and PDF release necessary for robust behavioural rhythmicity^38^. The s-LNv dorsal terminals exhibit circadian remodelling, with their terminals being more branched during the day than at night^40^ and having higher PDF accumulation in their terminals during the day than at night^38^.

The circuitry and molecular components of the mushroom body and the clock identified in *Drosophila* and shown to be highly conserved amongst insects^10,41,42^ making it a powerful model to test the effects of neonicotinoids on memory, circadian behaviour and sleep. They are also one of the insects whose nAChRs are best characterised. *Drosophila* have ten different nAChR subunits most of which are highly conserved across insect species, making it probable that a neurotoxin selected for its high potency to target insect nAChRs will affect the equivalent nAChR in beneficial insects^43^. Whilst the subunit conformation and location of neonicotinoid susceptible nAChRs is still largely unknown, in Drosophila the subunits Dα1 and Dβ2 have been shown to play a role in neonicotinoid susceptibility and resistance. Given the power of *Drosophila* as a model system, and the likely generalisation provided by conservation of nAChR function across insects, we tested the sub-lethal effect of field-relevant concentrations of the main banned and non-banned neonicotinoids on *Drosophila* memory, circadian rhythms and sleep.

## Results

Field relevant concentrations of neonicotinoids cause a range of lethal and sub-lethal effects in bees^1,18^. In order to validate the use of *Drosophila* as a model for these lethal and sub-lethal effects, we fed field relevant concentrations of the main banned and non-banned neonicotinoids to *Drosophila* and determined their effect on longevity, offspring viability and climbing ability. As in pollinators, longevity, fecundity and mobility were all affected by neonicotinoid exposure in Drosophila^11,44–46^. The mean lifespan of control flies was 49 days while exposure to field relevant concentration of 10 µg/L clothianidin causing a reduction to 28 days, imidacloprid and thiamethoxam to 36 days, and thiacloprid, which proved the least potent, to 39 days (Extended data Fig. 1). The viability of offspring was also reduced, with 100 µg/L clothianidin, thiamethoxam or thiacloprid and 10 or 100 µg/L imidacloprid reducing the proportion of eggs that subsequently completed development and eclosed as viable adults (Extended data Fig. 2). Likewise, field relevant concentrations of the banned neonicotinoids: imidacloprid, clothianidin and thiamethoxam (10 and 50 µg/L) all reduced locomotor performance, tested via a negative geotaxis climbing assay, whilst thiacloprid had no effect on locomotion at these concentrations (Extended data Fig. 3).

Olfactory associative memory is critical for foraging pollinators and has been shown to be disrupted by neonicotinoids in bees^47^. In order to see if field relevant concentrations of imidacloprid, clothianidin, thiamethoxam and thiacloprid had a similar effect on flies, one-hour (h) memory (Fig. 1) was assessed using *Drosophila* olfactory shock conditioning^48^. All three of the banned neonicotinoids reduced memory at 10 µg/L (Fig. 1a-c). The non-banned thiacloprid left memory intact (Fig. 1d). Sensory controls showed that none of the neonicotinoids tested reduced the ability of flies to sense either the odours or the aversive stimuli (Extended data Fig. 4). In order to localise the effect of nAChR mis-regulation on memory, we expressed a previously validated *RNAi* specific to the neonicotinoid susceptible nAChR subunits Dα1 and Dβ2^2^ throughout the mushroom body. Knock-down of either subunit was found to significantly reduce memory to a similar level as neonicotinoids (Fig. 1E), confirming the importance of these subunits in mushroom body mediated memory and suggesting the involvement of these subunits in the effect of neonicotinoids on this behaviour.

**Fig. 1.**
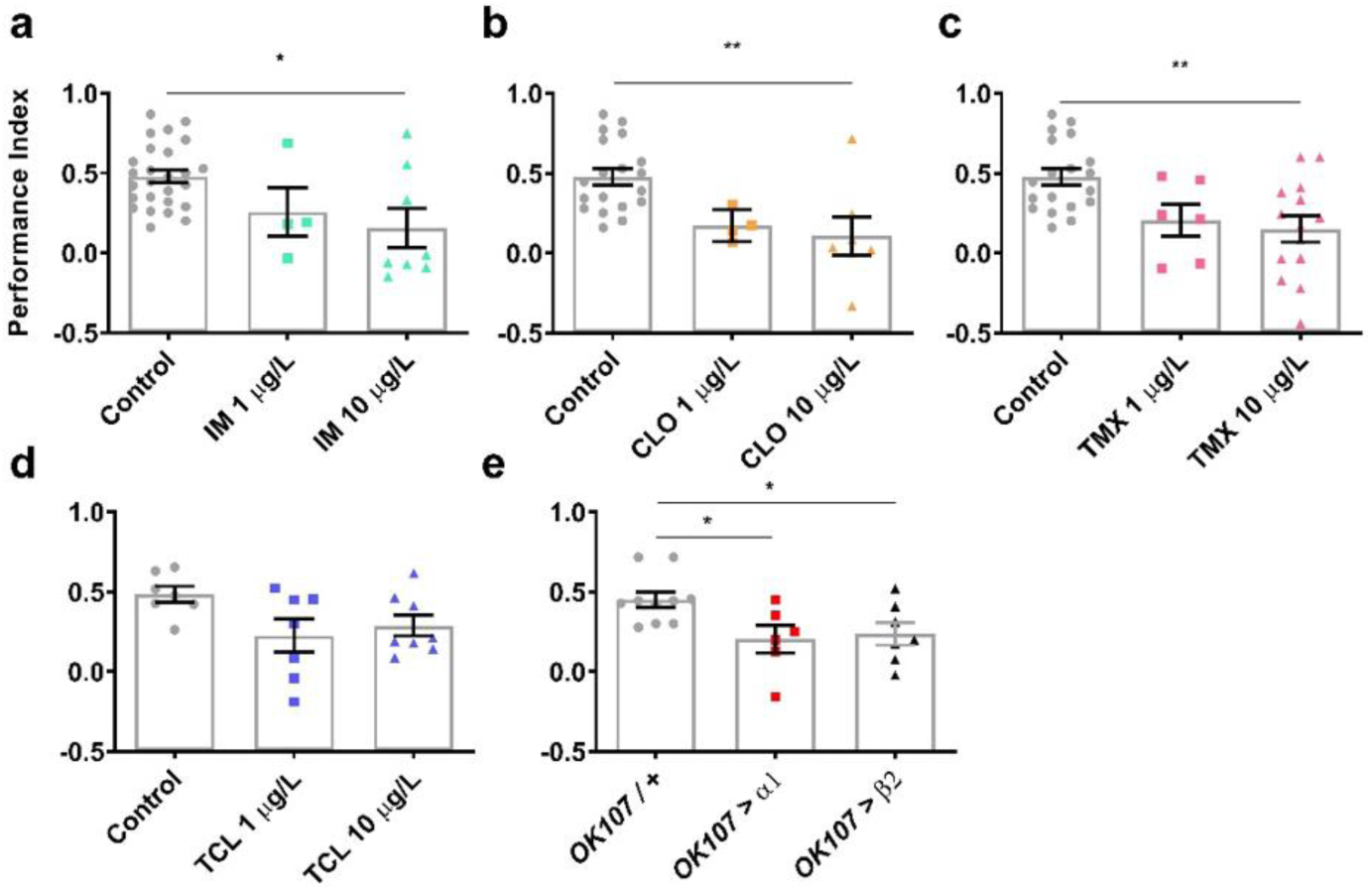
Field relevant concentrations of banned neonicotinoids or knockdown of Dα1 or Dβ2 in the mushroom bodies reduced 1 hour memory compared to control. 1h memory was reduced in flies exposed to field relevant concentrations of 1 or 10 μg/L of **a**, imidacloprid (IM) (*χ*^2^_2_)=7.3, *p*=0.026), **b**, clothianidin (CLO) (*χ*^2^_2_=12.4, p=0.002), **c**, thiamethoxam (TMX) (*χ*^2^_2_=9.6, p=0.008) and not in **d**, thiacloprid (TCL) (*χ*^2^_2_=5.0, p=0.084). **e**, Likewise, 1h memory was reduced in flies with *RNAi* mediated knockdown of Dα1 (*OK107-Gal4>uas-nAChR-Dα1*) or Dβ2 (*OK107-Gal4>uas-nAChR-Dβ2*) throughout the mushroom body (*F*_2,20_=4.6, *p*= 0.023). Each data point represents ~100 flies, n≥4 per treatment. Graphs show mean ± standard error of the mean (SEM) (*post hoc* pairwise comparisons: *p*≤0.05*, *p*≤0.01**, *p*≤ 0.001***, *p≤*0.0001****). The same tests, error bars and *p* values were used throughout.

These results extend existing data from pollinators showing a disruption of memory formation and showing for the first time that the disruption is mediated by nAChRs in the mushroom body. Similar investigation was then carried out on sleep and circadian rhythmicity, two other behaviours that are also heavily reliant on nAChR signalling. The effect of field relevant concentrations of neonicotinoids on circadian rhythms was tested using the *Drosophila* Activity Monitor (DAM2, Trikinetics Inc, USA)^49^. All three of the banned neonicotinoids caused a reduction of circadian rhythmicity (Fig. 2), with flies showing greatest sensitivity to the sub-lethal circadian effects of thiamethoxam, which caused a reduction in mean rhythmicity at 1, 10 and 50 μg/L (Fig. 2d) while clothianidin and imidacloprid both caused a reduction in mean rhythmicity at 50 μg/L (Fig. 2b, c). Any concentration tested of the three banned neonicotinoids caused an increase in the proportion of flies that were arrhythmic (rhythmicity statistic (R.S.) ≤1.5) compared to controls (Fig. 2f-I, Extended data Table 1). Again, the non-banned neonicotinoid, thiacloprid appeared not to have sub-lethal effects, with field-relevant concentrations of the insecticide leaving circadian rhythmicity intact (Fig. 2e, i).

**Fig. 2.**
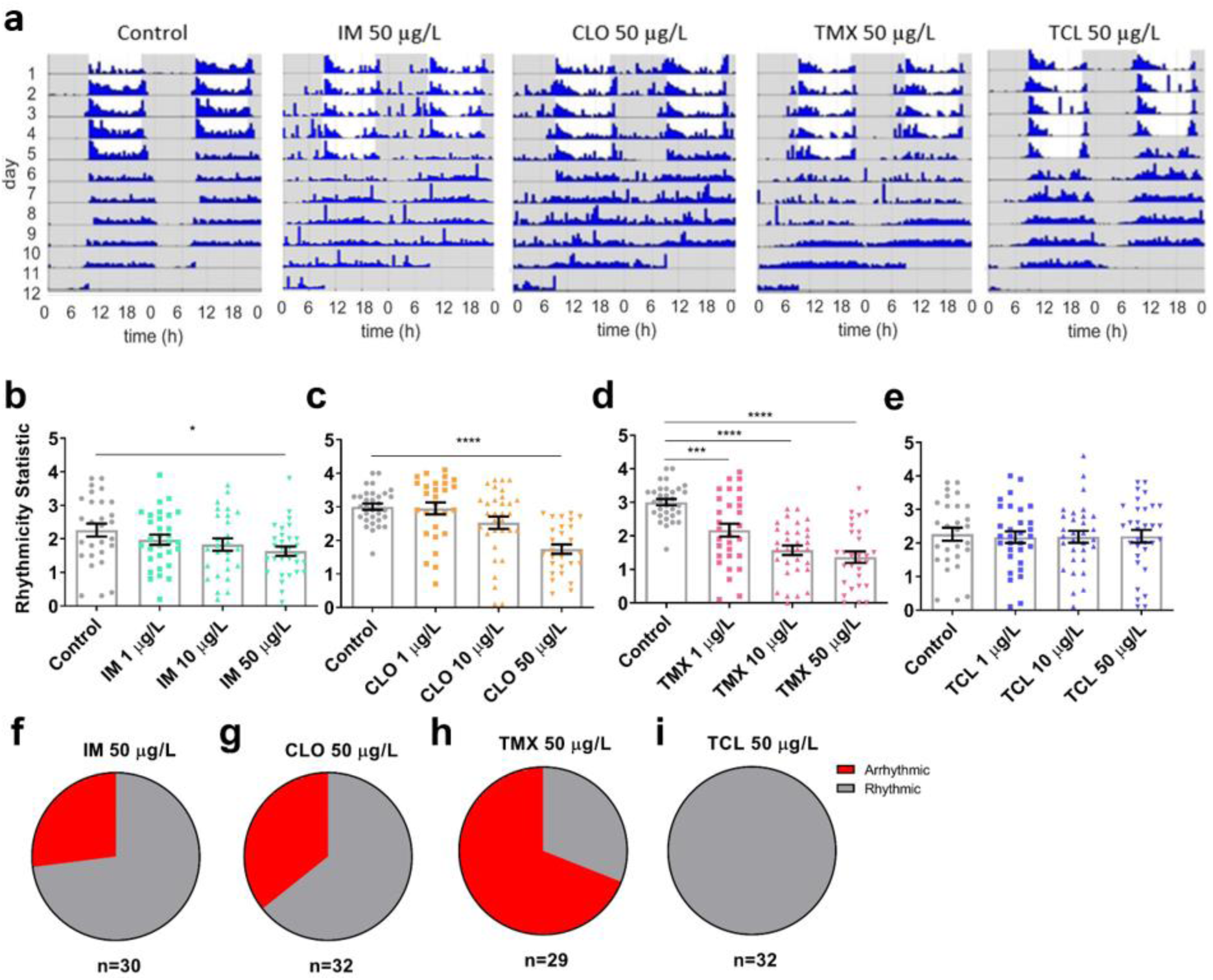
Field relevant concentrations of banned neonicotinoids reduced behavioural rhythmicity. **a**, Representative actograms of the activity of single flies for control or 50 μg/L of imidacloprd, clothianidin, thiamethoxam or thiacloprid. Mean rhythmicity for flies exposed to 1, 10 or 50 μg/L of **b**, IM (*F*_3,112_=2.5, *p*=0.06), **c**, CLO (*F*_3,116_=14.2, *p*<0.001), **d**, TMX (*F*_3,118_=23.7, p<0.001) and **e**, TCL (*F*_3,118_=0.05, p=0.987). Each data point represents a single fly, n=28-32 flies per treatment. Pie charts showing the increase in the proportion of the population who were arrhythmic (rhythmicity statistic (RS)≤1.5) for 50 μg/L of: **f**, IM, **g**, CLO, **h**, TMX and **i**, TCL, compared to controls.

Sleep was also monitored using the DAM system, with bouts of inactivity lasting more than 5 minutes qualifying as sleep^35^. Field relevant concentrations of all four neonicotinoids caused fragmentation of sleep, arising from sleep formed of a greater number of sleep episodes (Fig. 3b, e, h, l) of shorter length compared to control (Fig. 3c, f, I, m). This effect was greatest for clothianidin, where 1, 10 and 50 μg/L caused fragmentation of both daytime and night-time sleep (Fig. 3f, j) resulting in a reduction of night-time sleep (Fig. 3b). Thiamethoxam and imidacloprid had a similar effect (Fig. 3e, g, i, k) but only for night-time sleep (Fig. 3a, c). Thiacloprid caused an increase in the number of episodes initiated at night (Fig. 3h) and unlike the other neonicotinoids, caused a loss in daytime sleep (Fig. 3d) at every concentration tested, due to a reduction in daytime sleep episode length (Fig. 3l). This is likely due to the increase in daytime sleep latency observed in thiacloprid treated flies (Extended data Fig. 5).

**Fig. 3.**
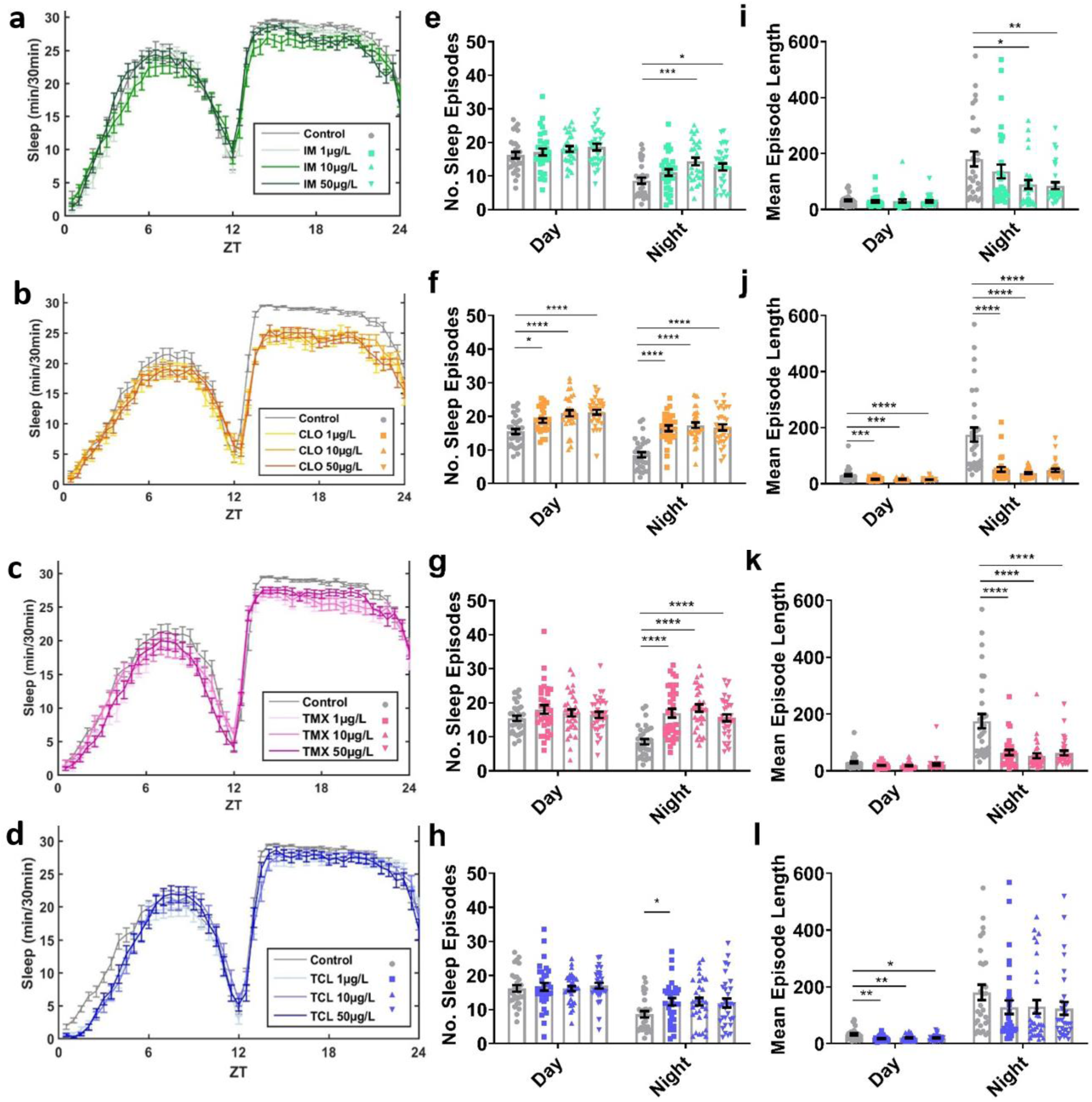
Field relevant concentrations of neonicotinoids disrupt sleep behaviour. Sleep plots showing the total sleep achieved per 30 minutes bin over the 24 h period (zeitgeber time (ZT)) for flies exposed to 1, 10 or 50 µg/L of **a**, imidacloprid, **b**, clothianidin, **c**, thiamethoxam or **d**, thiacloprid. The number of (no.) of sleep episodes initiated in **e**, IM, day (*F*_3,114_ = 1.2, *p* = 0.320) and night (*F*_3,114_ = 5.5, *p* = 0.001), **f**, CLO, day (*F*_3,120_ = 11.5, *p* < 0.001) and night (*F*_3,120_ = 25.0, *p* <0.001), **g**, TMX, day (*F*_3,124_ = 1.1, *p* = 0.344) and night (*F*_3,124_ = 17.0, *p* <0.001) or **h**, TCL, day (*F*_3,120_ = 0.2, *p* = 0.872) and night (*F*_3,120_ = 3.0, *p* = 0.034). Mean length (in minutes) of sleep episodes initiated in **i**, IM, day (*F*_3,114_ = 0.2, *p* = 0.889) and night (*F*_3,114_ = 4.5, *p* = 0.005), **j**, CLO, day (*F*_3,120_ = 9.9, *p* <0.001) and night (*F*_3,120_ = 21.8, *p* < 0.001), **k**, TMX, day (*F*_3,124_ = 2.5, *p* = 0.061) and night (*F*_3,124_ = 15.7, *p* < 0.001) or **l**, TCL, day (*F*_3,120_ = 5.2, *p* = 0.002) and night (*F*_3,120_ = 2.0, *p* = 0.121). Each data point represents a single fly, *n*=28-32 flies per treatment.

In order to localise the effects of neonicotinoids on sleep and circadian behaviour we specifically knocked down Dα1 or Dβ2 in all clock bearing cells. This resulted in reduced behavioural rhythmicity (Fig. 4b-d, g) and shorter night-time sleep episodes (Fig. 4f, i) in Dβ2 knock downs and caused sleep to be formed of a greater number of sleep episodes in both Dα1 and Dβ2 knockdown flies (Fig. 4e, h). This again showed that loss of these subunits caused similar behavioural disruptions as neonicotinoid exposure, suggesting a functional nAChR containing these subunits mediates the *in vivo* effects of these insecticides. In order to test whether this was the case, *RNAi* flies were exposed to 50 μg/L of imidacloprid or clothianidin, a concentration sufficient to reduce rhythmicity in control flies. On flies that already had their Dα1 or Dβ2 blocked genetically by expression of subunit specific *RNAi* expression throughout their clock, we found this caused no further loss of rhythmicity (Fig. 4j-k), providing evidence that the drug’s *in vivo* effects were mediated through a receptor containing one or both of these subunits in the clock.

**Fig. 4.**
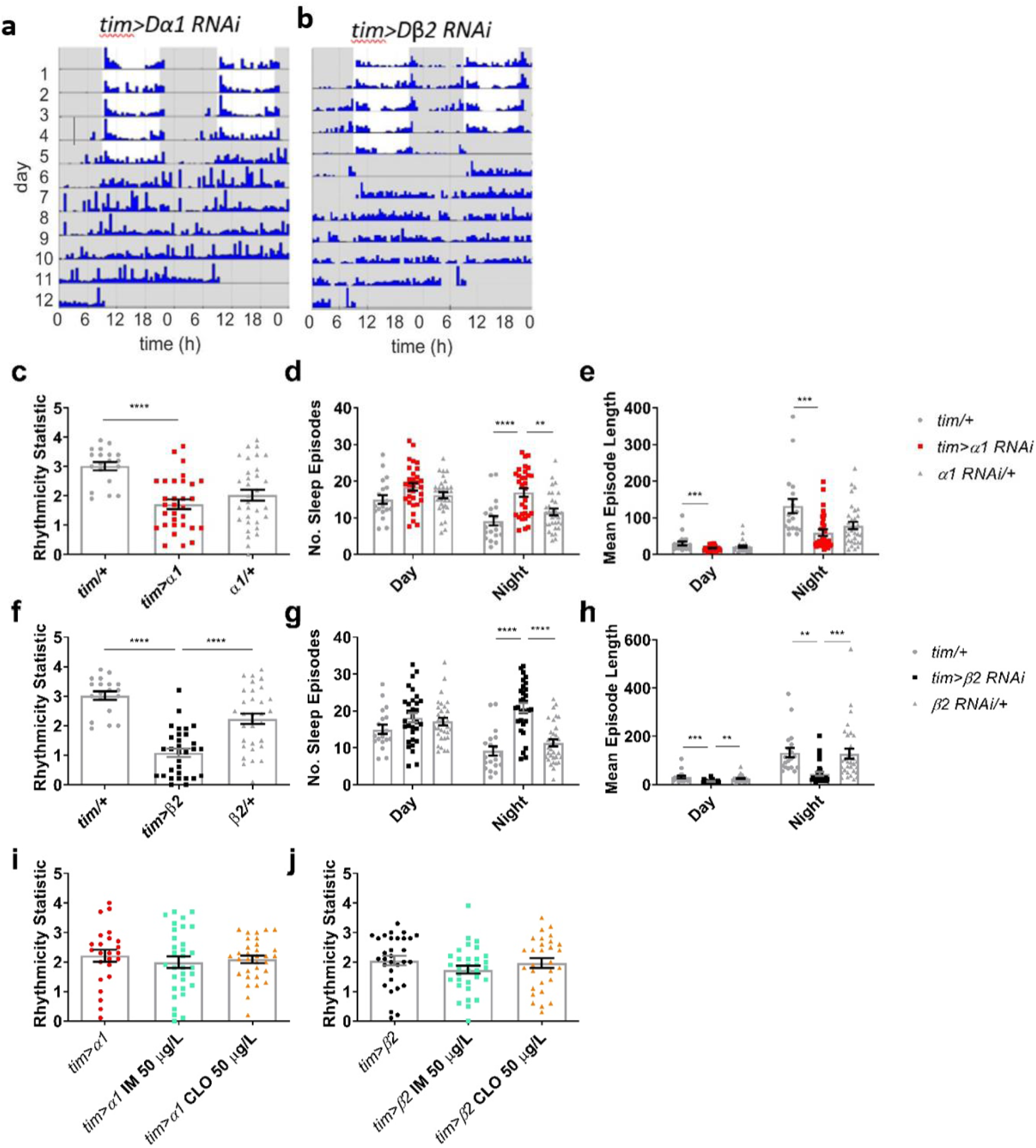
Knockdown of Dα1 or Dβ2 in the clock bearing cells disrupts circadian rhythmicity and sleep with no further effect by addition of neonicotinoids. Representative actograms for **a**, Dα1 knock down (*tim-Gal4>uas-nAChR-Dα1)* and **b**, Dβ2 knockdown (*tim-Gal4>uas-nAChR-Dβ2.* Effects of knocking down *Dα1* in clock bearing cells on **c**, rhythmicity (RS) (*F*2,79 = 11.8, *p* <0.001), **d**, number (no.) of sleep episodes in day (*F*_2,79_ = 2.9, *p* = 0.063) and night (*F*_2,79_ = 12.3, *p* <0.001) and **e**, mean episode length in day (*F*_2,79_ = 5.1, *p* = 0.008) and night (*F*_2,79_ = 8.3, *p* = 0.001). Effects of knocking down *Dβ2* in clock bearing cells on **f**, rhythmicity (*F*_2,79_ = 31.5, *p* <0.001), **g**, no. of sleep episodes in day (*F*_2,79_ = 1.6, *p* = 0.211) and night (*F*_2,79_ = 28.2, *p* <0.001) and **h**, mean episode length in day (*F*_2,79_ = 11.2, *p*<0.001) and night (*F*_2,79_ = 9.4, *p* <0.001). Each data point represents a single fly, *n*=19-32 flies per treatment. There was no additive effect of 50 µg/L of IM and CLO on **i**, *tim>α1* (*F*2,85 = 0.4, *p* = 0.677) and **j**, *tim>β2* (*F*_2,89_ = 1.1, *p* =0.336). Each data point represents a single fly, n=24-32 flies per treatment.

To further characterise the mechanism by which neonicotinoids disrupt circadian rhythms, the circadian remodelling and PDF cycling of the sLNv dorsal terminals were investigated. As previously reported^40^, in control flies the terminals were more branched and had higher accumulation of PDF in the day than at night (Fig. 5a-c).

**Fig. 5.**
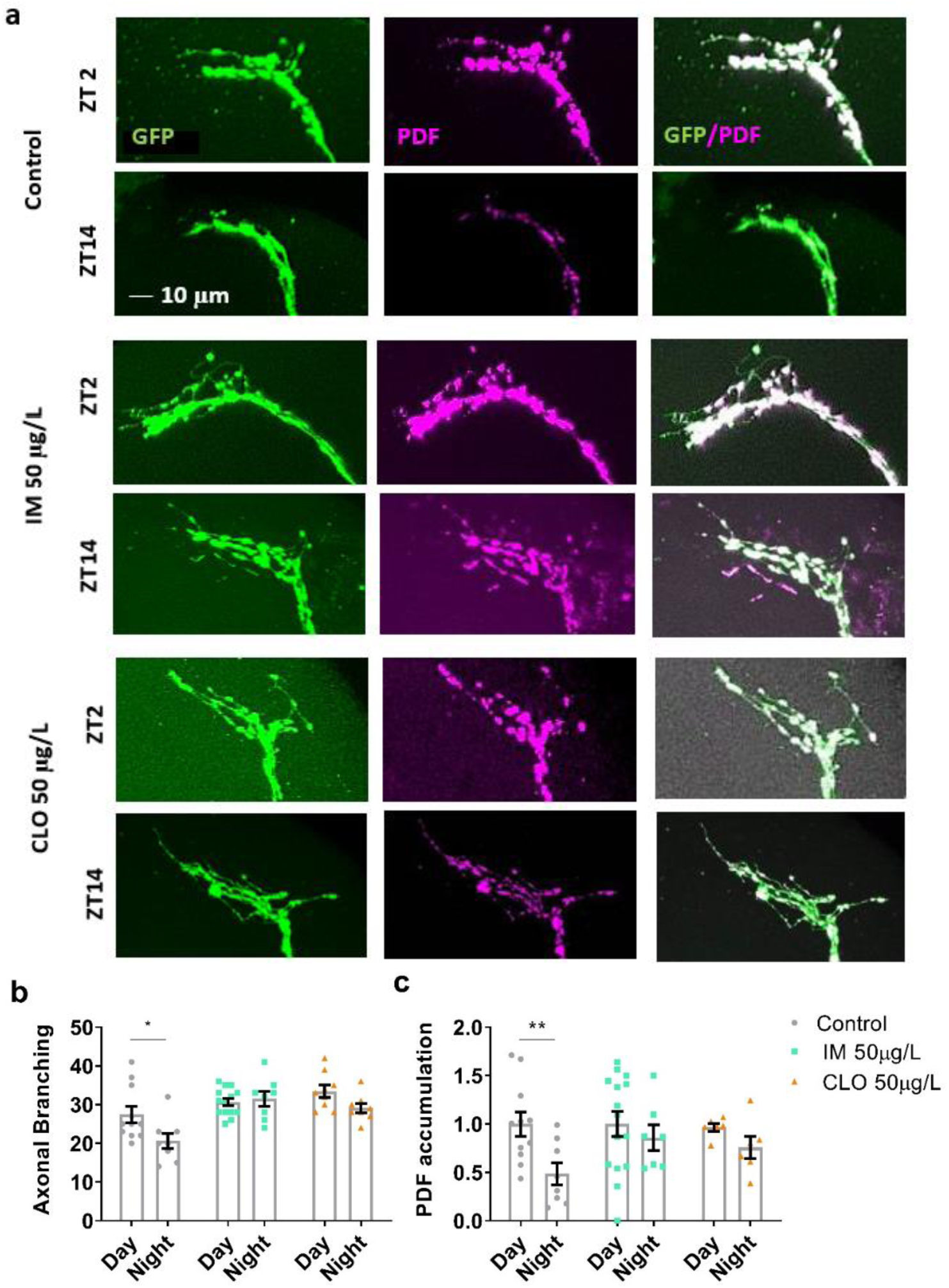
Field relevant concentrations of neonicotinoids disrupt the day/night remodelling and PDF cycling in the s-LNv clock neuron dorsal terminals. **a**, Representative confocal images of the s-LNv dorsal terminals for control and treated (50 µg/L IM or CLO) flies in the day (ZT2 i.e. 11am) and night (ZT14 i.e. 11pm). **b**, s-LNv dorsal terminal branching complexity is greater in the day than at night for control flies (*t*_17_=2.3, *p*=0.036). The day/night differences in complexity is removed in flies exposed to 50 µg/L of IM (*t*_14_=2.1, *p*=0.055) or CLO (*t*_15_=2.1, *p*=0.052). **c**, Accumulation of PDF in dorsal terminals is greater in the day than at night in control flies (*t*_17_=2.9, *p*=0.010), treatment with 50 µg/L IM (*t*_13_=1.0, *p*=0.332) or CLO (*t*_14_=2.1, *p*=0.054) removed this day/night difference in PDF levels. Each data point represents a single brain, n=6-15 brains.

In contrast, flies exposed to 50 μg/L imidacloprid or clothianidin showed no difference between day and night synaptic terminal branching or PDF accumulation (Fig. 5a-c). In flies with knockdown of either Dα1 or Dβ2 nAChRs in the PDF neurons, branching and PDF accumulation again showed no difference between day and night (Fig. 6a-c).

**Fig. 6.**
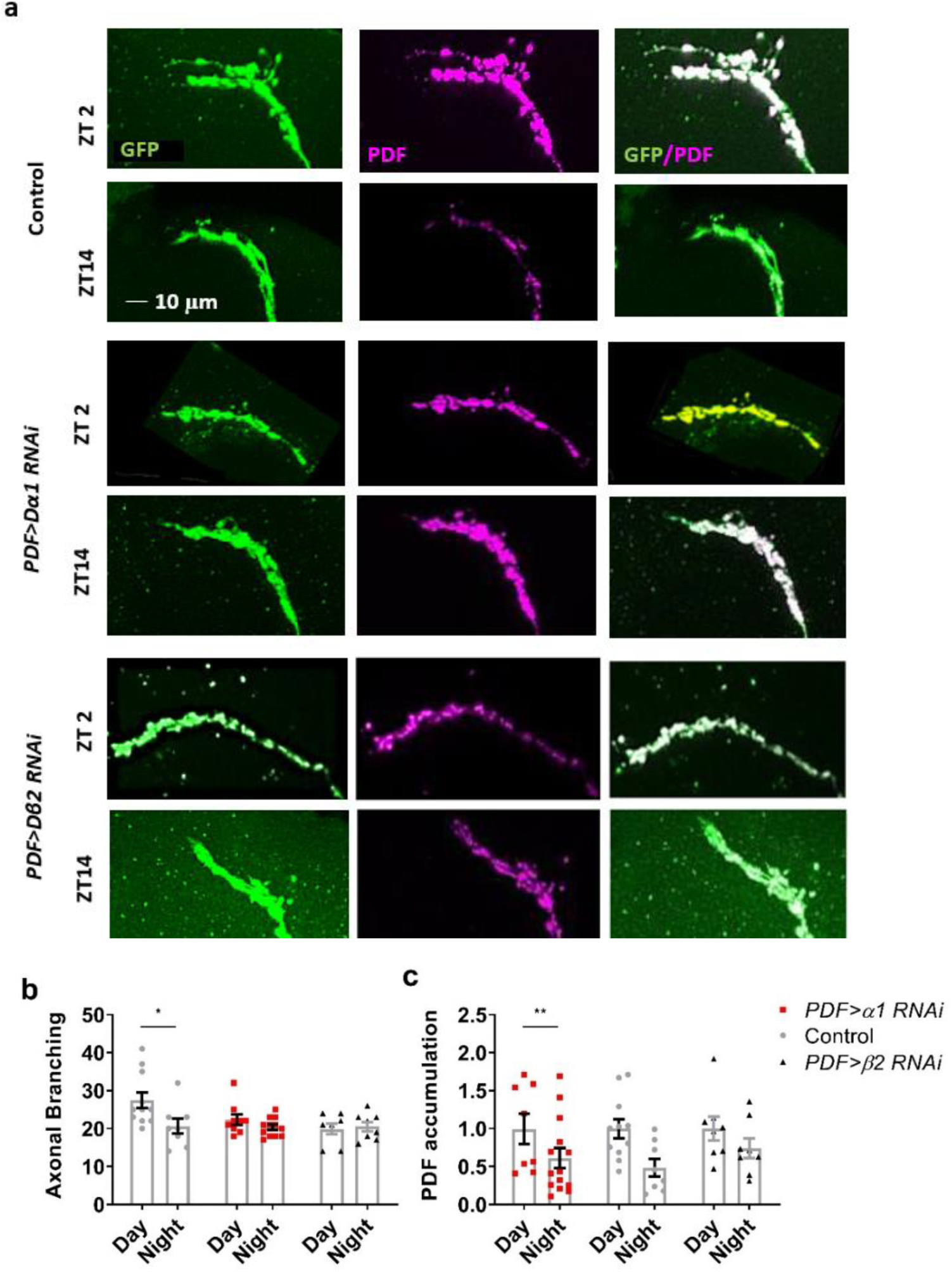
Knockdown of Dα1 or Dβ2 disrupted the day/night remodelling and PDF cycling of the s-LNv dorsal terminals. **a**, Representative confocal images of the s-LNv dorsal terminals of control flies and flies with Dα1 (*PDF-Gal4>uas-nAChR-Dα1-RNAi*) and Dβ2 (*PDF-Gal4>uas-nAChR-Dβ2-RNAi*) knocked down in LNv clock neurons taken in the day (ZT2) and night (ZT14). **b**, The s-LNv dorsal terminals of control flies showed greater branching complexity in the day than at night (*t*_17_=2.3, *p*=0.036), this day/night difference in terminal complexity was removed in *PDF>Dα1-RNAi* (*t*_19_=1.4, *p*=0.183) and *PDF>Dβ2-RNAi* (*t*_13_=-0.7, *p*=0.515) flies. **c**, PDF accumulation in the s-LNv dorsal terminals was greater in the day than at night for control flies (*t*_17_=2.9, *p*=0.010), but not in *PDF>Dα1-RNAi* flies (*t*_19_=1.8, *p*=0.089) and *PDF>Dβ2-RNAi* flies (*t*_14_=1.3, *p*=0.218). Each data point represents a single brain, n=6-15 brains.

## Discussion

Our data show that field relevant concentrations of all the neonicotinoids tested had some lethal effects in *Drosophila*, such as decreased viability and shortened lifespan. In contrast the behavioural or sub-lethal effects on flies differed between the banned and unbanned neonicotinoids with imidacloprid, clothianidin and thiamethoxam disrupting memory, locomotion, sleep and circadian behaviour, while the unbanned neonicotinoid thiacloprid only caused fragmentation and reduction in sleep, leaving the other behaviours intact. Thiacloprid appears to be less disruptive to the behaviours studied here than the three banned neonicotinoids. However its effects on sleep revealed significant sub-lethal effects even when exposure was at the lowest level reported (1 ppb), in addition to causing decreased viability and early death, providing strong evidence in support of the EU’s recommendation to use alternatives and perhaps to extend the neonicotinoid ban to thiacloprid. Clothianidin and thiamethoxam showed the greatest effects, which is consistent with them being full agonists at nAChRs and many other studies in pollinators finding them to be more toxic and potent than the partial agonist imidacloprid^1,17,50^.

The effects of field-relevant concentrations of neonicotinoids on memory in *Drosophila*, reiterates the conserved toxicity and sub-lethal effects of the insecticides on non-target insect species. That neonicotinoid exposure did not affect the ability of flies to sense the stimuli or respond to reinforcement, confirms that the neonicotinoids interfere with memory formation itself. This was supported by the data showing that knockdown of the neonicotinoid susceptible Dα1 or Dβ2 subunits in just the mushroom body was sufficient to cause the memory deficits observed in the neonicotinoid exposed flies. The Kenyon cells of the mushroom body are cholinergic^50^, with many of the projection neurons bringing olfactory information from the glomeruli of the antennal lobe forming nicotinic synapses onto the mushroom body^51^ and mushroom body to mushroom body output neuron synapses signalling via nAChRs^2^. Therefore, neonicotinoids can act at multiple nAChR synapses in the mushroom body circuit, disrupting the plasticity-relevant signals for memory formation. Furthermore in bees, field relevant concentrations of neonicotinoids disrupted mushroom body-mediated olfactory memory, electrically inactivated Kenyon cells^29^, decreased their synaptic density^30^ and reduced antennal lobe Ca^2+^ responses upstream of the mushroom body^31^.

Similarly, the sleep and circadian effects caused by neonicotinoid exposure appear to be due to the neonicotinoids acting directly upon the clock. Knockdown of Dβ2 nAChR subunit in the clock bearing cells resulted in the same disruption of rhythmicity as exposure to field-relevant concentrations of the banned neonicotinoids, whilst knockdown of Dα1 or Dβ2 caused changes to sleep behaviour reflecting those seen in neonicotinoid exposed flies. This suggests that Dα1 and Dβ2 mediate the effects of neonicotinoids on clock bearing cells, bringing about the disruptions in circadian rhythms and sleep. Consistent with this, exposure of Dβ2 knockdown flies to imidacloprid or clothianidin caused no further effect on circadian rhythmicity, confirming that Dβ2 in clock bearing cells mediates the *in vivo* effect of the banned neonicotinoids on circadian rhythms.

Given that the LNvs are nicotinic, neonicotinoids may be acting via these pacemaker neurons, which usually receive excitatory ACh inputs from light-sensing organs^3,4^. The electrical state of these neurons influences their circadian output including the circadian remodelling of the s-LNv dorsal terminals and circadian abundance of PDF^38,52^, with the release of this neuropeptide being necessary for behavioural rhythmicity^37^. We found that neonicotinoid exposure caused a loss of PDF cycling and terminal plasticity, with the terminals remaining in a branched, day state continuously, suggesting that the depolarising block effect of neonicotinoids can remove the normal day-night changes in the electrical state of the neurons required for circadian rhythms. Indeed, nAChR synaptic signalling is required for the rhythmic firing of action potentials in clock neurons in *Drosophila* and other insects^34,53,54^. Knock-down of the neonicotinoid susceptible nAChR subunits Dα1 or Dβ2 also removed day-night differences in the terminals as has been observed for flies with electrically silenced LNvs^52^. The agonist action of neonicotinoids on nAChRs on the LNvs may also explain the disruption to sleep behaviour. The electrical state of the l-LNvs is vital to their role as arousal neurons, with hyperexcitation of the l-LNvs leading to sleep defects such as loss of night-time sleep and shorter sleep episodes^55^ which we observed in the neonicotinoid exposed flies.

The high degree of structural and functional conservation of nAChRs between flies and bees^17,43,56^, and the conserved lethal and sub-lethal effects of reduced viability, longevity, locomotion and memory we demonstrated in flies as reported in bees suggest that the novel sleep and circadian disruptions we observed are also likely to occur in beneficial insects in the field^11,44–47^. Previous work has shown that sub-lethal effects observed in the lab translate to the field, for example neonicotinoid reduced foraging motivation observed for bumblebees both in the lab and in free flying colonies in the field^28,57^. Reduced behavioural rhythmicity is likely to reduce the amount of activity carried out in the daytime, reducing pollination and foraging opportunities^9^. The reduction in total sleep and fragmentation of sleep will reduce the quantity of deep sleep achieved, as deep sleep occurs later into the sleep episode^58^. As deep sleep is particularly important for memory consolidation^8^, this may compound the direct effects of neonicotinoids on memory, which will again impact foraging efficiency.

In summary, nAChR subunits Dα1 and Dβ2 expression in the mushroom body appears to mediate the effect of field-relevant concentrations of the banned neonicotinoids on memory, while expression of these subunits in clock bearing cells mediates the effect of the banned neonicotinoids on circadian rhythmicity and sleep. The non-banned neonicotinoid, thiacloprid, was less toxic than the banned insecticides, only disrupting sleep, which seems to be the most sensitive behavioural metric of the sub-lethal effects of neonicotinoids. In addition, both the banned and unbanned neonicotinoids decreased both viability and shortened lifespan, therefore supporting the EU recommendation to seek alternatives to their use as well as supporting their continued ban. This work illustrates the utility of neonicotinoids as a pharmacological tool for exploration of nAChR function, as well as the use of *Drosophila* in revealing the mechanism of action of neonicotinoids and elucidating the sub-lethal and lethal effects of these insecticides, highlighting their potential impact on insects in the field.

## Methods

### Fly husbandry and genotypes

The following fly stocks were used: wild-type strains *iso31* (Gift from Dr Ralph Stanewsky, University of Münster, Germany) for climbing, circadian and sleep assays and *CSw-*(gift from Dr Scott Waddell, University of Oxford, UK) for all other experiments, *Pdf-Gal4* (Bloomington *Drosophila* stock center number (BDSC): 6900), *elav-Gal4* (BDSC: 8760), *tim-Gal4*[27] (gift were from Dr Ralf Stanewsky, University of Münster, Germany), *OK107-Gal4* (BDSC: 854), *uas-mcd8::GFP* (BDSC: 5137), *uas-nAChR-Dα1-RNAi* (BDSC: 28688), and *UAS-nAChR-Dβ2-RNAi* (BDSC: 28038) validated in^2^. For all experiments, flies were collected shortly after eclosion and used within 5 days. For climbing, circadian, sleep, longevity, immunohistochemistry and offspring survival assays females were used, for learning and memory mixed sex groups were used.

Flies were bred, maintained and tested on standard polenta-based food mixture at 25°C, 55-65% humidity under 12h light:12h dark (LD) conditions. Food was made up in 5 L quantities and contained: 400 g polenta, 35 g granulated agar, 90 g active dried yeast, 50 g soya flour, 400 ml malt extract and 200 ml molasses, with 40 ml of propionic acid (Sigma-Aldrich, #94425) and 100 ml of nipagin (Sigma-Aldrich, #H5501) added once cool. Neonicotinoids were added to food before it set from a frozen and aliquoted stock solution of 100,000 μg/L ddH_2_O. The neonicotinoids were analytical standard (PESTANAL Sigma-Aldrich): imidacloprid, clothianidin, thiamethoxam, and thiacloprid.

### Longevity

Ten once mated, one day old females were placed in a vial containing control or neonicotinoid containing food and transferred into a fresh vial every 2 days with the number of dead flies noted. This was continued until all flies were dead^59^ with ten repeats being performed per treatment group. A survival curve was created and analysed using GraphPad (GraphPad Prism version 6.05 for Windows, GraphPad Software) and mean lifespan calculated. The difference of the treatment survival curve from the control survival curve was analysed using a log-rank (Mantel-Cox) test.

### Offspring viability

Flies were reared on control or neonicotinoid containing food. Ten groups of ten once mated females were collected, and then each female was placed in a vial of control fly food and allowed to lay eggs over a 24hour period. The number of eggs in each vial was quantified and then compared to the number of adult flies which successfully eclosed from the vial ~15 days later, giving a % offspring survival for each group^60^.

### Locomotor assay

Climbing ability was used as a measure of locomotion of adult flies and was determined by the negative geotaxis assay, whereby flies were tapped to the bottom of a tube, causing them to move away from gravity (negative geotaxis). Twenty-five groups of ten females were placed in vials of control or neonicotinoid containing food for 5 days. They were then placed into empty vials. After 5 minutes of acclimatisation, flies were knocked to the bottom of the vial and given 10 seconds to climb^61^. The performance index represents the proportion of flies who successfully climbed ≥7 cm in 10 seconds.

### Aversive olfactory conditioning

1h memory was tested using aversive olfactory conditioning^48^. Groups of 30-50 one-five days old mixed sex flies were reared on control or neonicotinoid containing food. Flies were exposed consecutively to one of two odours, either 4-methylcyclohexanol (Sigma) or 3-octanol (Sigma) diluted 1:500 and 1:250 respectively, paired with 1.5 second pulses of 70 V electric shock, with 3.5 second pauses between shocks. Flies were then returned to food vials for 1 hour before memory was tested. For testing, flies were loaded into the choice point of the T-maze and, after a 90 second acclimatisation period, were given the choice of two tubes, one containing each of the test odours. Flies were given 2 minutes and then the proportion in each arm was counted. A separate group of flies were then trained and tested with the reciprocal odour. The performance index score represents the proportion of flies who correctly avoided the arm containing the odour which had been delivered with shock during training, as shown below. PI = (number of correct flies−number of incorrect flies)/total number of flies.

The PI for flies shocked with each odour separately were averaged to give an *n*=1.

### Sensory Controls

Sensory controls were carried out to check the capacity of treatment groups to sense olfactory and shock cues^48^. For olfactory acuity, groups of 1-5 day old 30-50 mixed sex flies, reared on control or neonicotinoid food, were loaded into the T-maze and provided with a choice between an odour (1:500 4-methylcyclohexanol or 1:250 3-octanol) and fresh air. For shock reactivity, similar groups of flies were given a choice between two shock tubes, one of which was delivering 1.5 second pulses of 70 V electric shock, with 3.5 second pauses between shocks. In both cases, flies with normal sensory capacity should avoid the stimuli. The proportion of each group who avoided the odour or shock was reported.

### Circadian rhythms

Behavioural rhythmicity data was collected using *Drosophila* Activity Monitors (DAM2, TriKinetics Inc)^62^. Virgin females were placed in individual tubes in the DAM, with control or neonicotinoid containing food, 32 flies per treatment, for 5 days in LD followed by 5 days constant darkness (DD). Flies who died before the end of day ten were not included in analysis. DAM tubes were intersected with an infrared beam, with each beam cross counted as an activity bout.

To quantify circadian rhythmicity, the data was summed into 30 minutes bins. From this an actogram was created and rhythmicity statistic and period length were calculated for each individual fly for the DD portion using Flytoolbox^63^ in MATLAB (MATLAB and Statistics Toolbox Release 2012b, The MathWorks). The proportion of flies in each treatment group whose Rhythmicity Statistic was below 1.5, generally considered to denote arrhythmicity^63^, was calculated. This was then displayed in a pie chart for each group, after being normalised by removing the proportion of controls who were arrhythmic for each run.

### Sleep

For analysis of sleep behaviour, flies were loaded into the DAM as described above. Sleep measures were extracted from activity data from 5 days of LD, summed into one minute and 30 minutes bins. Sleep was defined as bouts of inactivity lasting more than 5 minutes, as is convention^63,64^. The mean total sleep, mean sleep episode length, mean number of sleep episodes per day and night and mean sleep latency were calculated for each individual using the Sleep and Circadian Analysis MATLAB Program (SCAMP) in MATLAB^65^. For each treatment, a mean sleep profile was also plotted showing mean sleep quantity per 30 minutes bin over the 24 h period.

### Measuring the arborisation and PDF cycling of s-LNv dorsal terminals

Immunohistochemistry was adapted from the method described in Fernández et al^40^. Virgin females were collected and placed in vials. After 5 days LD, on control or neonicotinoid food, flies were anesthetised using CO2 at either 2h after lights on (ZT2) or 2h after lights off (ZT14) and decapitated. Heads were fixed in phosphate-buffered solution (PBS) with 4% paraformaldehyde (Thermo Fisher Scientific) containing 0.008% Triton X-100 (Sigma-Aldrich) for 45 minutes at room temperature. Heads were washed quickly twice in PBS with 0.5% Triton X-100 (PBT 0.5%), followed by three 20 minutes washes in PBT 0.5% and dissection in PBT 0.1%. Brains were blocked in 5% Normal Goat Serum (NGS, Thermo Fisher) for 1h, then incubated for 36 hours at 4ºC with mouse monoclonal anti-PDF (Developmental Studies Hybridoma Bank, #PDF-C7) and rabbit polyclonal anti-GFP (Life Technologies # A11122) in NGS at concentrations of 1:200 and 1:1000 respectively.

Brains were washed again as before and then incubated for 3h at room temperature followed by 24h at 4 °C with Alexa Fluor Plus 488 Goat anti-mouse (Life Technologies # A32723) and Alexa Fluor Plus 555 goat anti-rabbit (Life Technologies # A32732) in NGS at concentrations of 1:1000 and 1:100 respectively. Brains were washed once more and then mounted onto glass slides using spacers (SecureSeal™, Grace Bio-Labs #654002), covered with VectaShield hard set medium (Vector Laboratories) and secured with CoverGrip (Biotium #23005).

Imaging was carried out on a Leica SPE confocal laser scanning microscope with the green channel imaged at 480–551 nm and the red at 571–650 nm. Z stacks were captured of the s-LNv dorsal terminals using a 64x oil immersion objective, with a step size of 2 µm. Maximal projection stacks were created and analysed using FIJI (ImageJ) ^66^. The arborisation of each s-LNv terminal was calculated using an adaptation of the Scholl analysis^40^. Six concentric rings 10 μm apart were drawn, centred at the first branching point and the number of branches touching each ring was summed. Both hemispheres were measured. Mean scores for ZT2 and ZT14 in each group were compared using Pearson’s *t*-test using SPSS Statistics 24 (IBM Corporation).

For PDF staining intensity, the image was cut at the first branching point to create an image containing only the terminals and not the cell axon and the image was analysed in FIJI (ImageJ). The image was transformed into 8-bit and the threshold adjusted to create a black and white image. The despeckle filter was used to reduce noise and watershed segmentation carried out to separate the different PDF compartments. This image was then used as a template for calculating the PDF staining in the original maximal projection image, allowing the PDF staining intensity to be calculated for each of these compartments and the mean taken. The mean for both hemispheres of the brain was calculated and reported, and the means for ZT2 and ZT14 were compared as for axonal branching.

### Statistical analysis

Normality of the data was checked using a Shapiro-Wilk test. Data were also checked for homogeneity of variance using Levene’s test for equality of variances. Unless otherwise stated, means were then compared using a one-way ANOVA with *post hoc* pairwise comparisons being carried out using Tukey’s multiple comparisons test. Where data were not normally distributed, a Kruskal-Wallis one-way ANOVA was carried out with Dunnett’s multiple comparison. Statistical analysis was done in SPSS Statistics 24 (IBM Corporation). Graphs were created in GraphPad (Prism version 8.0.0 for Windows, GraphPad Software).

The sleep data failed the assumptions of normal distribution and homogeneity of differences. Thus, permutation tests were conducted in R 3.4.1. As the resulting *p* values closely matched those produced by analysing the same data using a one-way ANOVA as above, and because ANOVA is relatively robust to deviations from normality when sample sizes are large, the results of the one-way ANOVA were displayed.

## Acknowledgements

We thank Drs Ralph Stanewsky, University of Münster, Germany and Scott Waddell, University of Oxford, UK for providing flies. We thank Drs X.X. for providing comments on the manuscript and acknowledge the Wolfson Bioimaging facilities at University of Bristol. This work was supported by a BBSRC studentship BB/J014400/1 and Leverhulme project grant RPG-2016-318 awarded to J.J.L.H.

## Author Contributions

J.J.L.H., K.T. and S.A.R. designed the study. K.T. performed and analysed the circadian and sleep assays, immunohistochemistry work, climbing and longevity and lethality assays. S.H. carried out the learning and memory assays. E.B. performed the electrophysiological recordings. K.T. and J.J.L.H. wrote the paper with input from S.A.R., E.B. and S.H. The project was supervised by J.J.L.H. and S.A.R., who secured funding and edited the manuscript.

