## Extended Data for "Neonicotinoids disrupt memory, circadian behaviour and sleep"

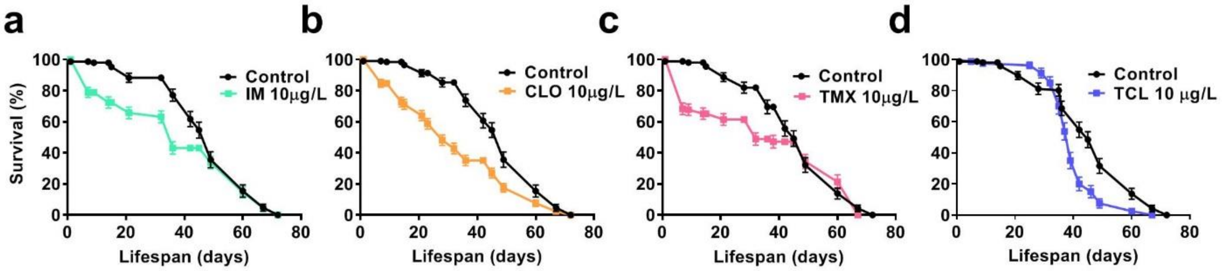


**Extended data Fig. 1| Field relevant concentrations of neonicotinoids reduced longevity.** Compared to the median lifespan for control flies (49 days) flies exposed to 10 µg/L of **a,** IM (36 days,( $\chi^2_1$=8.3, *p*=0.004), **b,** CLO (28 days, ($\chi^2_1$=41.0, *p* <0.001), **c,** TMX (36 days, ($\chi^2_1$=5.8, *p*=0.016) and **d,** TCL (39 days, ($\chi^2_1$=18.5, *p* <0.001) had shorter lives. *n*=100 flies for each group.

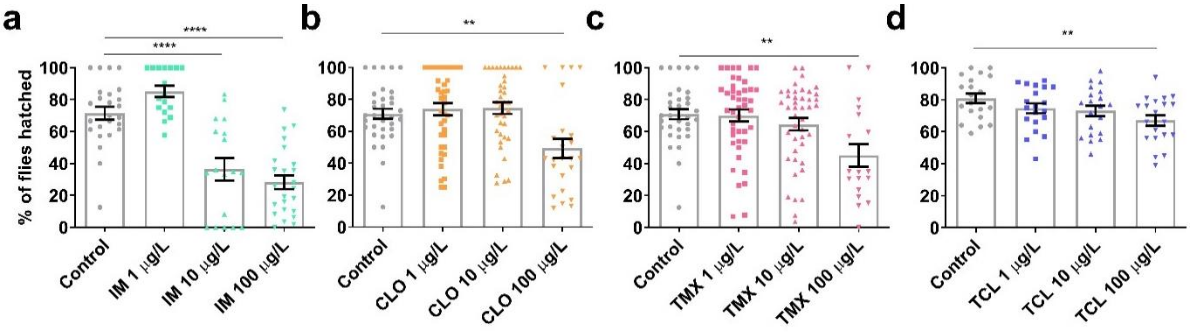


**Extended data Fig. 2| Field-relevant concentrations of neonicotinoids reduce viability.** The viability of offspring of flies exposed to 1, 10 or 100 µg/L of **a**, IM ($F_{3,80}$ = 31.7, $p \leq$ 0.001), **b,** CLO ($F_{3,132}$ = 6.9, *p* <0.001), **c,** TMX ($F_{3,135}$ = 5.4, *p* = 0.002) and **d**, TCL ($F_{3,76}$ = 3.8, *p* = 0.014), *n*=10 groups of 10 once mated female flies for each treatment. Viability was measured by counting the number of eggs laid by 10 once mated female in 24 h period and then counting the % of those eggs that completed development, eclosing viable adults ~15-18 days later.

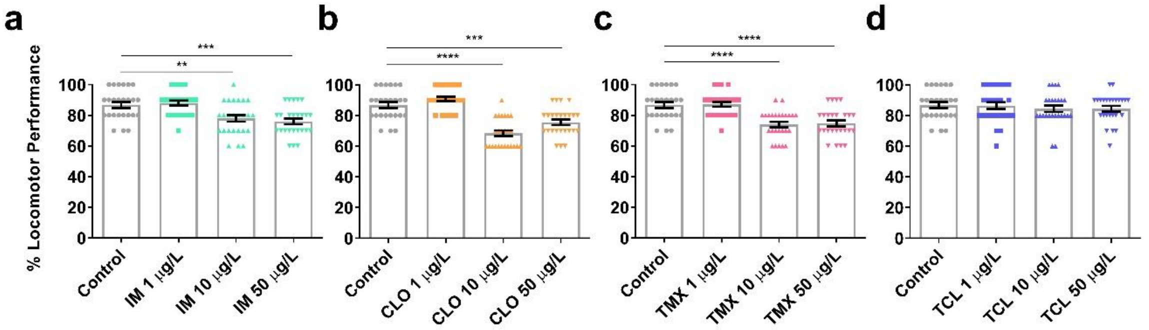


**Extended data Fig. 3| Field relevant concentrations of neonicotinoids reduce locomotor performance.** Locomotor performance was measured using the negative geotaxis climbing assay, flies exposed to 10 or 50 µg/L of the banned neonicotinoids: **a,** IM ($F_{3,96}$ = 9.9, $p$ <0.001), **b,** CLO ($F_{3,96}$ = 32.0, $p$ <0.001), **c,** TMX ($F_{3,96}$ = 15.8, $p$ <0.001) significantly reduced climbing performance, while the non-banned neonicotinoid **D)** TCL ($F_{3,96}$ = 0.4, $p$ = 0.762) did not affect locomotion. $n$=25 flies for each treatment group.

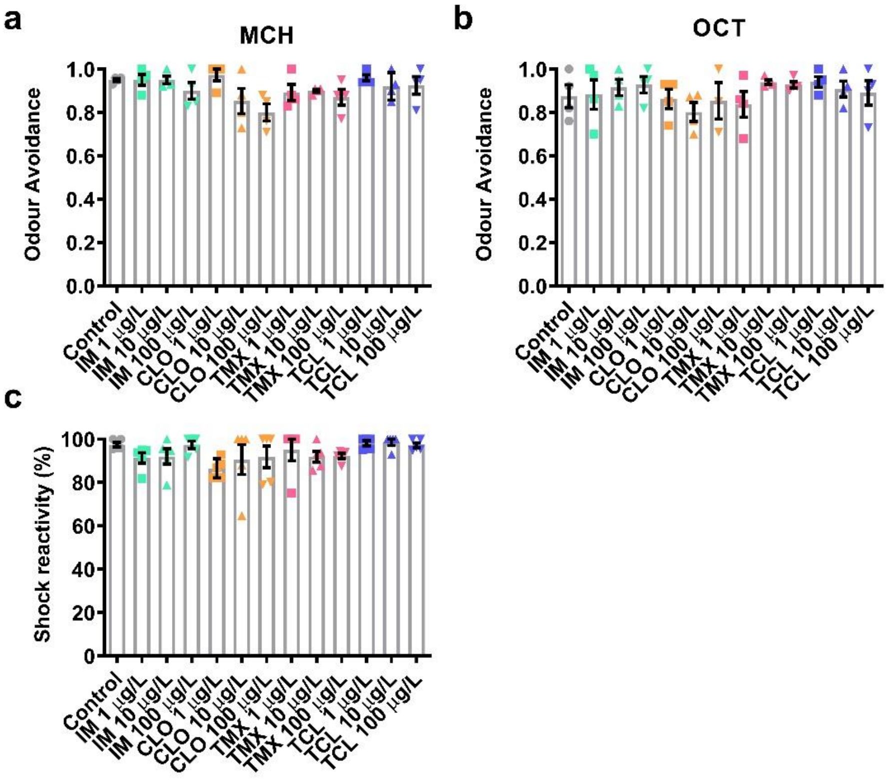


**Extended data Fig. 4| Field relevant concentrations of neonicotinoids do not disrupt olfaction or shock reactivity.** Sensory controls for olfactory-shock conditioning memory assays (Fig. 1) show 1, 10 and 100 µg/L of IM, CLO, TMX and TCL did not affect **a,** odour avoidance of 4-methylcyclohexanol (MCH) ($\chi^2_{12}$ = 19.5, $p$ = 0.076), or **b,** 3-octanol (OCT) ($\chi^2_{12}$ = 10.0, $p$ = 0.674) and **c,** shock reactivity ($\chi^2_{12}$ = 22.6, $p$ = 0.031). Each data point represents a group of ~50 flies, tested together, $n$=4 for each group.

**Extended data Table 1| Field relevant concentrations of neonicotinoids increase the proportion of the population (%) exhibiting arrhythmicity compared to controls**

| | IM | CLO | TMX | TCL |
|---|---|---|---|---|
| 1 µg/L | 8% | 11% | 10% | 1% |
| 10 µg/L | 19% | 19% | 23% | 4% |
| 50 µg/L | 27% | 36% | 65% | 0% |

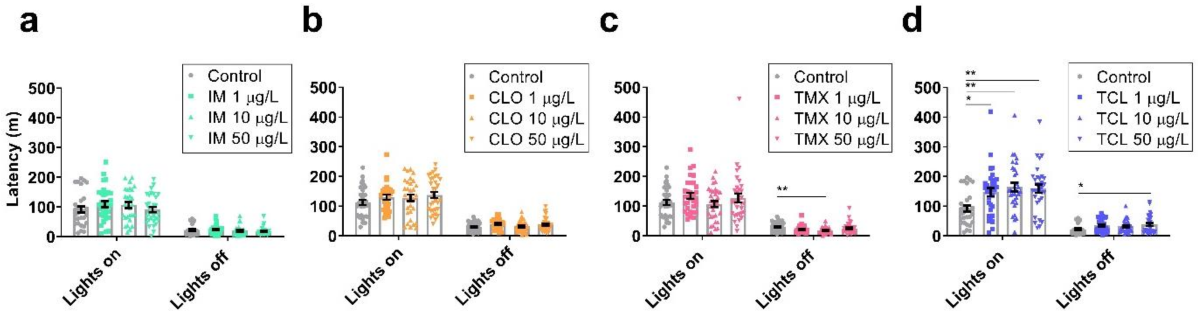


**Extended data Fig. 5| Field relevant concentrations of thiacloprid increase daytime sleep latency.** The mean latency in minutes (m) before sleep was initiated after lights on or lights off, for flies exposed to 1, 10 or 50 µg/L of **a,** IM, day ($F_{3,114}$=0.9, p=0.441) and night ($F_{3,114}$=0.9, p=0.468), **b,** CLO, day ($F_{3,120}$=1.1, p=0.333) and night ($F_{3,120}$=2.0, p=0.124), **c,** TMX, day ($F_{3,124}$=1.3, p=0.264) and night ($F_{3,124}$=4.3, p=0.007) or **d,** TCL, day ($F_{3,120}$=5.6, p=0.001) and night ($F_{3,120}$=2.7, p=0.50). Each datapoint represents a single fly, n=28-32 flies per treatment.
